# Mitotic spindle formation in the absence of Polo kinase

**DOI:** 10.1101/2021.12.15.472863

**Authors:** Juyoung Kim, Gohta Goshima

## Abstract

Mitosis is a fundamental process in every eukaryote, in which chromosomes are segregated into two daughter cells by the action of the microtubule (MT)-based spindle. Despite this common principle, genes essential for mitosis are variable among organisms. This indicates that the loss of essential genes or bypass-of-essentiality (BOE) occurred multiple times during evolution. While many BOE relationships have been recently revealed experimentally, the BOE of mitosis regulators (BOE-M) has been scarcely reported and how this occurs remains largely unknown. Here, by mutagenesis and subsequent evolutionary repair experiments, we isolated viable fission yeast strains that lacked the entire coding region of Polo-like kinase (Plk), a versatile essential mitotic kinase. The BOE of Plk was enabled by specific mutations in the downstream machinery, including the MT-nucleating γ-tubulin complex, and more surprisingly, through downregulation of glucose uptake, which is not readily connected to mitosis. The latter bypass was dependent on casein kinase I (CK1), which has not been considered as a major mitotic regulator. Our genetic and phenotypic data suggest that CK1 constitutes an alternative mechanism of MT nucleation, which is normally dominated by Plk. A similar relationship was observed in a human colon cancer cell line. Thus, our study shows that BOE-M can be achieved by simple genetic or environmental changes, consistent with the occurrence of BOE-M during evolution. Furthermore, the identification of BOE-M constitutes a powerful means to uncover a hitherto under-studied mechanism driving mitosis and also hints at the limitations and solutions for selecting chemotherapeutic compounds targeting mitosis.

## Introduction

Different organisms have different sets of essential genes for their viability and propagation (1, 2). This indicates that most ‘essential’ genes are context-dependent and can become dispensable during evolution. Plausibly, a loss of essentiality is compensated for by manifestation of an alternative, currently ‘masked’, or far less active mechanism to ensure a similar cellular activity. Many experimental efforts have been made to recapitulate the molecular diversity found in nature (3). Large-scale systematic surveys have been recently conducted in budding and fission yeasts, in which a number of bypass-of-essentiality (BOE) events have been identified (4–7). In these studies, suppressors were screened for haploid strains in which an essential gene was experimentally disrupted. For 9–27% of the essential gene disruptants, a mutation or overexpression of other gene(s), or chromosomal gain makes the strain viable, indicating that essentiality depends on genetic background and that BOE could indeed occur at a certain frequency. However, in most cases, the underlying mechanism remains unexplored. It is also unclear why BOE is rarely observed in certain processes such as mitotic cell division.

Mitotic cell division is controlled by many essential genes in a given cell type (8–10). Evolutionary evidence of BOE is clearly visible for this fundamentally critical biological process. One striking example is the centrosome, which is assembled by the action of many essential proteins in animal, fungal, and algal species, and plays a vital role in cell division and cellular motility (11). However, the centrosome and most of its components have been lost in land plants, and yet plant cells undergo spindle assembly and chromosome segregation at high fidelity (12). Kinetochore components, such as the CCAN complex, spindle microtubule (MT)-associated proteins, such as TPX2, augmin, and mitotic motors, and cell cycle regulators, such as anaphase-promoting complex/cyclosome are among other examples; they are not universally conserved or essential factors (12–15). Despite the evidence of BOE for almost all the genes involved in mitosis, only a limited number of cases can be found in experimental BOE screening. For example, two random BOE screenings in fission yeast encompassing 23 mitotic genes have identified only a single protein, Cnp20/CENP-T, despite the fact that >20% BOE has been observed for mitosis-unrelated genes (6, 7). The BOE of *cnp20* is conceivable, as CENP-T functions in parallel with CENP-C for kinetochore assembly (16, 17). Another known BOE of mitosis regulators (BOE-M) in fission yeast is MT plus-end-directed kinesin-5/Cut7, which is required for bipolar spindle formation through force generation on spindle MTs. The viability of *cut7*Δ was restored when the opposing minus-end-directed kinesin-14/Pkl1 was simultaneously deleted. Thus, the balance of forces applied to spindle MTs is critical (18–20). However, many other essential mitotic genes have no apparent functionally redundant or counteracting factors, and whether these relationships are general mechanisms of BOE-M is unclear. Essential mitotic genes are potential targets of cancer chemotherapy (21); it is also important to understand the BOE that underlies drug resistance.

In this study, we found that the essentiality of the sole Polo-like kinase (Plk) in fission yeast (Plo1) can be bypassed. Plo1, similar to human Plk1, is assumed to be essential for spindle MT formation and spindle bipolarisation. However, these essential processes were restored in the absence of Plo1 by multiple independent mechanisms that increase MT nucleation and stabilisation, one of which involved a remarkably simple change in glucose concentration in the culture medium and depended on casein kinase I (CK1). Thus, our study uncovered an unexpected alternative mechanism of spindle MT formation and further implies that more BOE-M can be recapitulated in the experimental system.

## Results

### Viable yeast cells without Polo-like kinases in several genetic backgrounds

Our previous BOE screening randomly selected 93 genes on chromosome II, which encompassed 12 mitotic genes (7). BOE-M could not be detected in any of these genes. To further screen for BOE-M, we selected eight other mitotic genes (*ark1, bir1, fta2, fta3, mis6, mis14, pic1*, and *plo1*) and applied the same screening method. Seven days after plating and UV mutagenesis of spores of each disruptant, we found a growing haploid colony for *plo1*Δ, the sole Polo-like kinase (Plk) in *Schizosaccharomyces pombe* (Fig. 1A, first step). Plks play versatile roles in animal cell division, including centriole duplication (by Plk4), centrosome maturation, spindle assembly checkpoint satisfaction, and cytokinesis (by Plk1). It is also a possible target for cancer chemotherapy (21–23). The responsible suppressor mutation was identified through whole genome sequencing (WGS) followed by genetic crossing, which turned out to be *ght5* (Fig. 1B, left, fourth line). In *S. pombe*, eight hexose transporters have been identified, which show different affinities to glucose; Ght5 is a major hexose transporter with the strongest affinity to glucose and plays a critical role in glucose uptake (24). This prompted us to test and found that the *plo1*Δ strain grows, albeit slower than the wild-type, when the glucose concentration of the medium was lower than 0.3% (Fig. 1B, left, third line, Fig. S1A). Thus, *plo1*Δ became viable under low-glucose conditions. To obtain a full scope of suppressor mutations, we repeated the UV mutagenesis of *plo1*Δ spores on a larger scale, obtained multiple colonies, and determined the responsible mutations. Simultaneously, an ‘experimental evolution’ (EVO) (also called ‘evolutionary repair’ (3)) experiment was conducted for the *plo1*Δ strain in low-glucose medium (0.08%), in which serial dilution and saturation enriches the cells that have acquired beneficial mutations for proliferation (Fig. 1A, ‘1^st^ EVO’). The faster growing strains obtained through this step were subjected to further evolutionary repair experiments in high-glucose medium (‘2^nd^ EVO’) and at a different temperature (‘3^rd^ EVO’).

**Figure 1.**
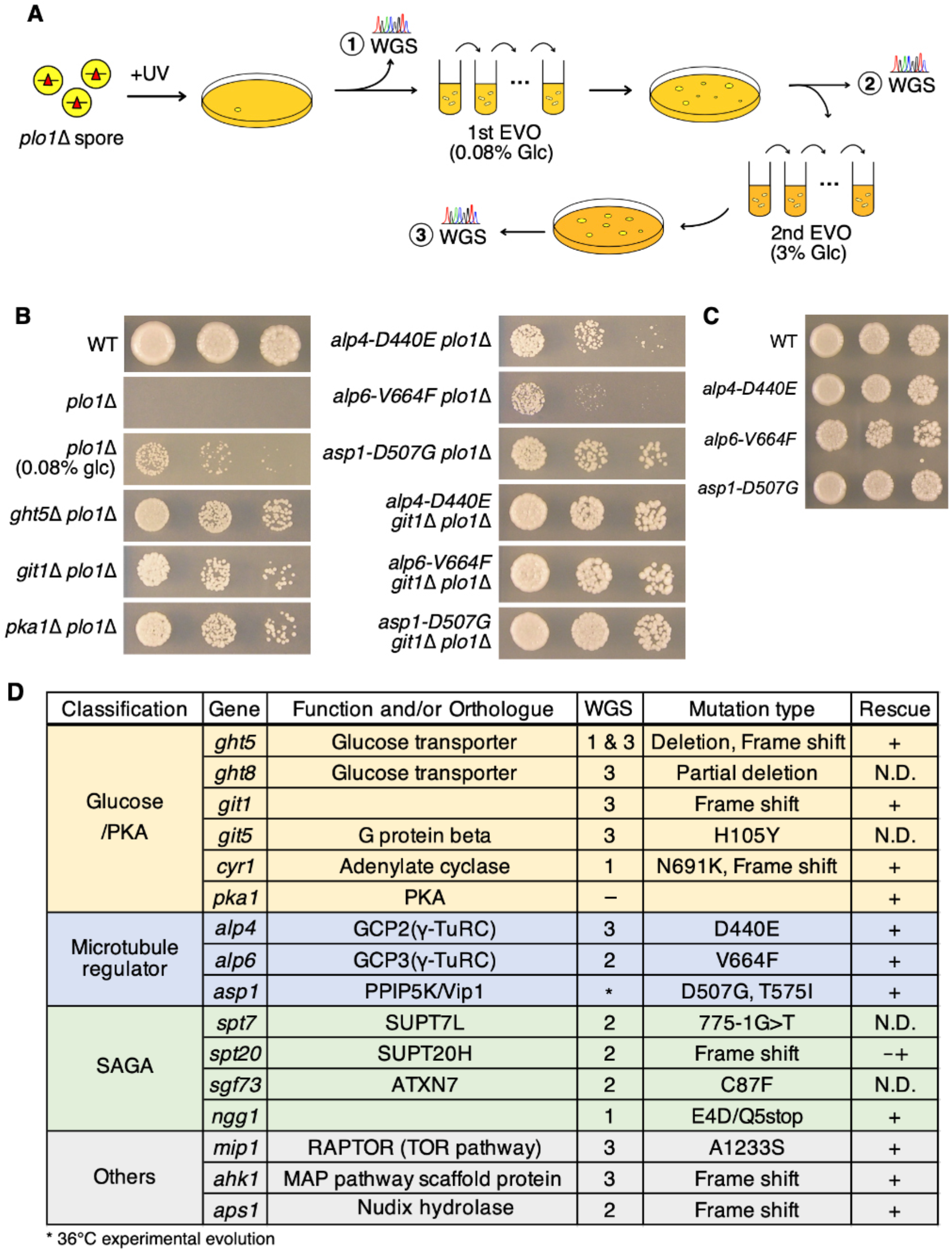
Isolation of viable *plo1*Δ strains. (A) Experimental procedure to isolate *plo1*Δ strains. The yeast spores in which *plo1* gene was replaced by drug-resistant cassette were mutagenised by UV and plated onto the drug-containing medium. The haploid colonies that appeared after several days represent the *plo1*Δ strains. The whole genome sequence (‘WGS’) was determined to map the responsible suppressor mutations, while a few strains were subjected to experimental evolution (EVO), where serial dilution and saturation accumulated fitness-increasing mutations. EVO was repeated three times in different conditions, and suppressor mutations were determined by WGS. (B) Viable *plo1*Δ strains obtained by indicated suppressor mutations. Cells (5,000, 1,000, and 200) were spotted onto normal YE5S plates, except for the third row, where glucose (glc) concentration in the medium was reduced to 0.08% (YE5S, 4 d, 32°C). (C) Single mutants of *alp4-D440E, alp6-V664F*, and *asp1-D507G*. Cells (5,000, 1,000, and 200) were spotted onto normal YE5S plates and incubated for 3 d at 32°C. (D) List of suppressor mutations for *plo1*Δ. The ‘WGS’ column indicates at which step in (A) the mutation was identified. ‘Rescue’ column indicates whether the indicated mutation alone bypassed the essentiality of Plo1. The colony grew extremely poorly for the *spt20*Δ *plo1*Δ (marked with -+).

We determined the whole genome sequences of several viable *plo1*Δ strains and confirmed the suppressor mutations by independently generating a double mutant with *plo1*Δ. In total, 16 genes were found to assist in the growth of the otherwise inviable *plo1*Δ strain (Fig. 1B, D; S1B, C). An example of the evolutionary repair process is shown in Fig. S1D. This strain acquired mutations in *alp6* and *aps1* during the 1^st^ EVO but still possessed the benefit of *plo1*^+^ gene for strain fitness (Fig. S1D, left). However, additional mutations in *mip1* and *ahk1* during the 2^nd^ EVO bypassed the requirement of Plo1, since adding back the *plo1*+ gene to the original locus did not further promote colony growth (Fig. S1D, right). Most of the responsible genes were categorised into three classes: Spt-Ada-Gcn5 acetyltransferase (SAGA) complex, glucose/PKA pathway, and MT regulators (Fig. 1D). The SAGA complex is a general regulator of transcription, possessing histone acetyltransferase activity, and affects the expression of many genes (25). We did not analyse this in the present study. The cAMP-PKA pathway is linked to glucose homeostasis in fission yeast. Glucose is detected by a receptor (Git3), and the G-protein complex (Gpa2, Git5, and Git11) is activated, which then activates adenylate cyclase (Cyr1) to produce cAMP (26). Eventually, cAMP releases the inhibitor Cgs1 from Pka1, converting Pka1 to its active form (27). Furthermore, yeast cells regulate glucose uptake by changing the localisation and transcriptional level of hexose transporters, including Ght5, depending on environmental conditions (24, 28, 29). There is a link between glucose/PKA and MT stabiliser CLASP (cytoplasmic linker-associated protein) during interphase (30). In our case, mutations in MT regulators (*alp4, alp6*, and *asp1*) and glucose/PKA-pathway genes additively supported the growth of *plo1*Δ (Fig. 1B).

### Monopolar spindles predominate during mitosis in the absence of Polo

The major MT nucleator at the centrosome is the γ-tubulin ring complex (γ-TuRC), which consists of γ-tubulin and GCP subunits, including GCP2 (Alp4/Spc97) and GCP3 (Alp6/Spc98) (31). In animal cells, Plk1 is a critical regulator of mitosis, which is, in the early stage, required for γ-TuRC recruitment to the centrosome and thus centrosome maturation; inhibition of Plk1 leads to monopolar spindle formation (22, 32). In fission yeast, cytokinesis/septation defects are most profoundly observed in *plo1* mutants, whereas monopolar spindle formation has also been described (33–37). However, actual spindle dynamics have not been analysed for *plo1*Δ in live imaging. To analyse spindle dynamics in the absence of Plo1, live imaging of mCherry-tubulin and a spindle pole body (SPB) marker, either Sad1^SUN^-GFP or Alp6^GCP3^-GFP, was performed after *plo1*Δ spore germination with spinning-disc confocal microscopy (Fig. 2A-E). The control cell assembled bipolar spindles immediately after the disappearance of interphase MT networks (Fig. 2A and D: time 0 corresponds to the onset of mitosis), and cell division was completed in ~30 min. In contrast, 53 out of 56 *plo1*Δ cells after spore germination were arrested with a monopolar spindle for >1 h (Fig. 2B, C, E, wherein stronger laser exposure was applied in Fig. 2C to visualise the faint MT signals).

**Figure 2.**
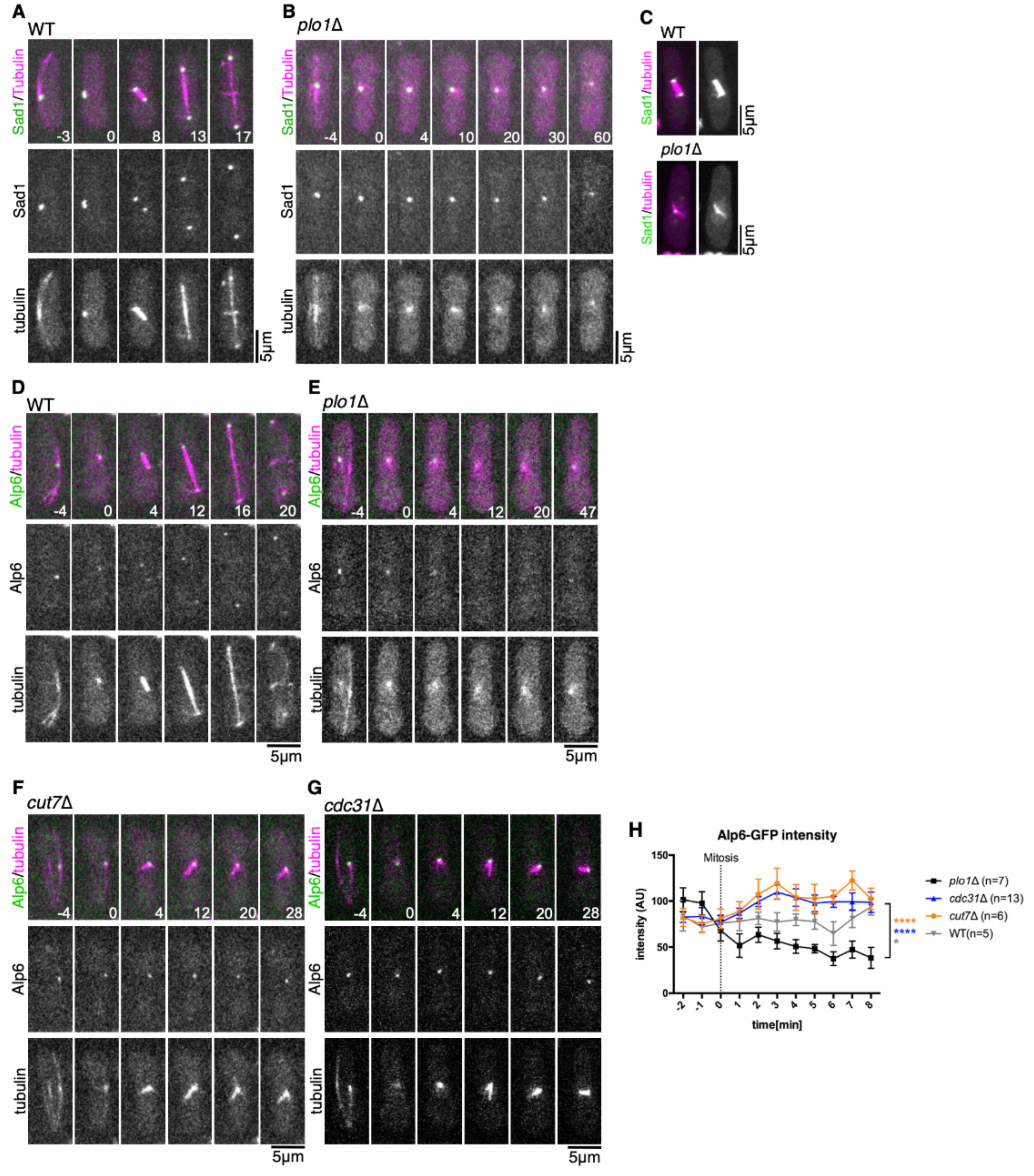
Monopolar spindle formation with reduced microtubules (MTs) and γ-TuRC localisation in *plo1*Δ. (A, B) Live imaging of the control and *plo1*Δ strains expressing Sadl^SUN^-GFP and mCherry-tubulin. The first mitotic phase after spore germination was imaged. (C) Mitotic spindles of control and *plo1*Δ strains expressing Sadl^SUN^ -GFP and mCherry-tubulin with longer exposure time. (D, E) Live imaging of the control and *plo1*Δ strains expressing Alp6^GCP3^-GFP and mCherry-tubulin. (F, G) Monopolar spindles of the *cut7*Δ and *cdc31*Δ. strains after germination. (H) Quantification of Alp6^GCP3^-GFP intensity during mitosis. The signal intensities from 5 to 8 min were compared between the *plo1*Δ and wild-type (WT), *cut7*Δ, or *cdc31*Δ (****, p < 0.0001; *, p = 0.0226). Error bars indicate the standard error of the mean (SEM). Time 0 (min) was set at the onset of spindle formation.

We compared the phenotype with other known mutants that show monopolar spindle formation, including Cut12, which drives SPB insertion into the nuclear envelope (NE) (38, 39), Cut7^kinesin-5^, which is required for anti-parallel MT crosslinking and sliding (40), and Cdc31^centrin^, which is required for SPB duplication (41). In the *cut12-1* temperature-sensitive (ts) mutant, the lack of insertion of one of the duplicated SPBs causes partial breakage of the NE and detachment of an SPB from the NE (38, 42). In another study, Plo1 was shown to regulate the formation of the Sad1^SUN^ ring structure, which might be required for SPB insertion (43). We assessed the integrity of the NE in *plo1*Δ. First, we tagged GFP to Pcp1^PCNT^, a core SPB component (35, 44), in the *plo1*Δ background. In contrast to the *cut12-1* mutant, we always detected punctate Pcp^PCNT^-GFP signals (n = 20) at the pole of the monopolar spindle, suggesting that SPBs in the NE generate spindle MTs in the absence of Plo1 (Fig. S2A, B). Next, we conducted an NLS-GFP efflux assay, in which partial NE breakage because of SPB insertion error leads to nuclear GFP signal efflux into the cytoplasm (38). We first confirmed the efflux in the *cut12-1* ts mutant; GFP started to leak out from the nucleus 18 ± 10 min after interphase spindle disassembly at non-permissive temperatures in 20 out of 20 cells that assembled monopolar spindles (Fig. S2D, 20, 30, and 70 min; (38)). In contrast, *plo1*Δ cells maintained GFP signals inside the nucleus during the early stage of mitosis, similar to the control strains (Fig. S2C). These data suggest that SPBs are properly inserted into the NE at the onset of spindle formation in the absence of Plo1. Notably, GFP efflux during mitotic arrest occurred in 16 out of 20 *plo1*Δ cells (69 ± 18 min after spindle assembly), suggesting that NE was partially dissolved during prolonged arrest (Fig. S2C, 100 and 200 min).

Next, we isolated *cut7*Δ and *cdc31*Δ spores with Alp6^GCP3^-GFP (SPB) and mCherry-tubulin markers and germinated them in normal culture medium. As expected, monopolar spindles were prevalent in each sample, with only a single dot of Alp6^GCP3^-GFP detectable at the end of spindle MTs (Fig. 2F, G). However, Alp6^GCP3^-GFP signal intensity was significantly lower in the *plo1*Δ spindles than in *cut7*Δ or *cdc31*Δ (Fig. 2H). Consistent with this phenotype, spindle MTs were dimmer in *plo1*Δ (compare Fig. 2E and 2F/G) and *plo1*Δ was sensitive to thiabendazole (TBZ), an MT-destabilising drug (Fig. S3A). Finally, we checked whether Cut7^kinesm-5^ localisation was defective in *plo1*Δ, which would cause spindle monopolarisation. Cut7^kinesin-5^-GFP accumulation at the SPB and spindle was delayed in the absence of *plo1* (Fig. S3B, 0 min). However, the signals gradually recovered and reached a level comparable to the early prometaphase of control cells, at which spindle bipolarity was not recovered (Fig. S3C). Thus, it is unlikely that failure in Cut7^kinesin-5^ recruitment is the major cause of spindle monopolisation in *plo1*Δ. Rather, our data favour the model whereby decreased MT nucleation at SPB leads to spindle monopolisation in *plo1*Δ. Consistent with this notion, monopolar spindles have been observed in mutants of the γ-TuRC component (Alp4^GCP2^) (45).

### Modulation of MT nucleation and stability bypassed Polo essentiality

Two BOE strains had point mutations in *alp4^GCP2^* and *alp6^GCP3^*. Double *alp4-D440E plo1*Δ and *alp6-V664F plo1*Δ strains recovered colony-formation ability in normal (high-glucose) medium (Fig. 1B). Thus, the essentiality of Plo1 was bypassed by a single specific mutation in the MT nucleating machinery. We also performed a spot test for single *alp6-V664F* and *alp4-D440E* mutants (Fig. 1C). *alp6-V664F* grew more slowly than the wild-type, whereas no difference in colony growth was observed for *alp4-D440E*.

We investigated whether the mutation in *alp4* could restore γ-TuRC recruitment to the SPB in the absence of *plo1*. To address this, we isolated a double *alp4-D440E plo1*Δ mutant with Alp6^GCP3^-GFP and mCherry-tubulin markers (Fig. 3A). Quantification indicated that both GFP and mCherry signals were partially but significantly restored by the *alp4-D440E* mutation (Fig. 3B, C). In 50% of the cells (n = 46), spindle bipolarity was recovered after a delay and cytokinesis was completed (Fig. 3A, left), whereas monopolar states were persistent for >60 min in 30% of the cells (Fig. 3B, right), explaining the partial rescue of the viability by this specific mutant of *alp4*. Consistent with frequent spindle bipolarisation, GFP efflux in the viable *alp4-D440E plo1*Δ and *ght5*Δ *plo1*Δ strains was less frequently observed than in single *plo1*Δ (19% and 21%, n = 26 and 48) (Fig. S2E, F).

**Figure 3.**
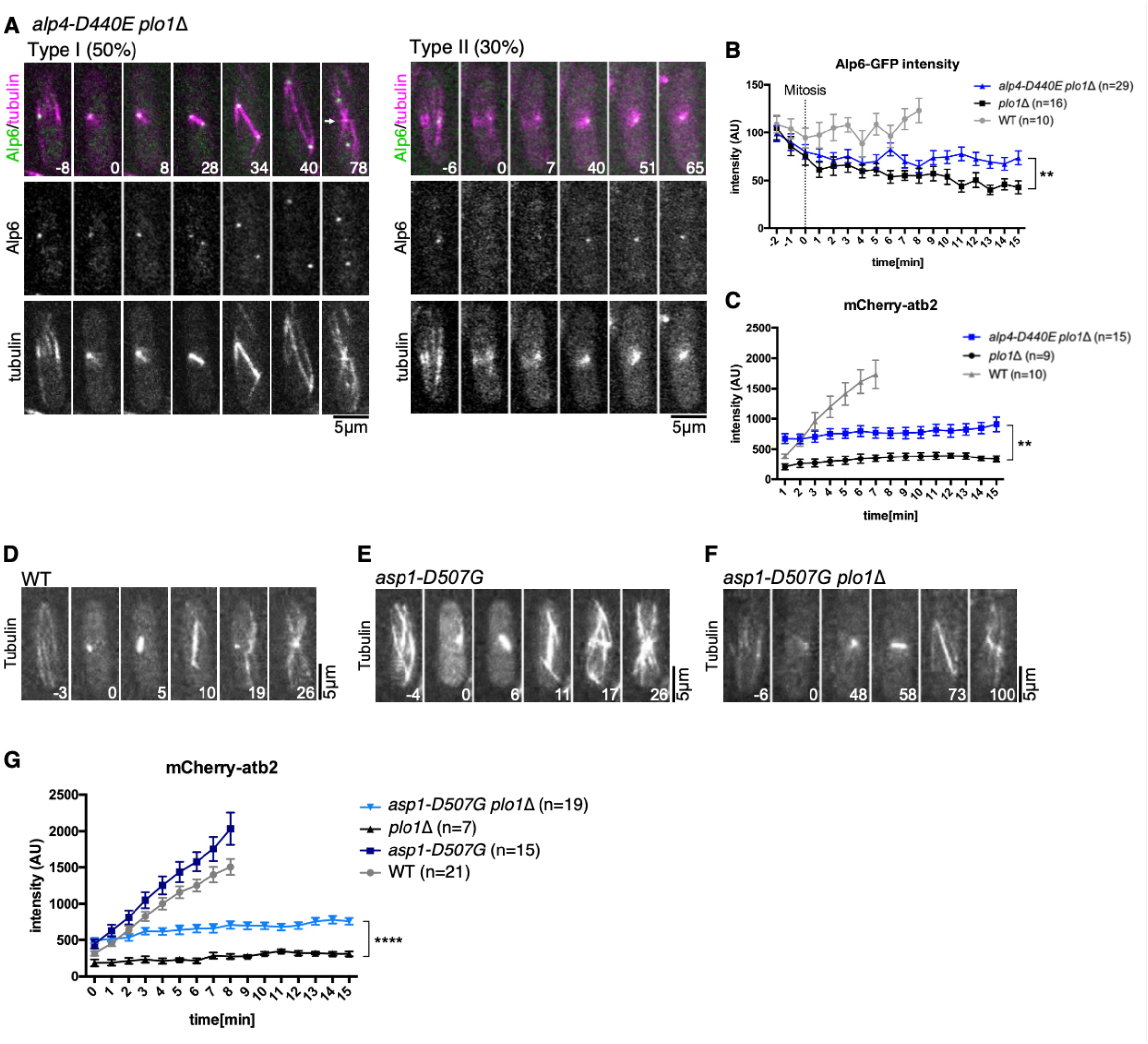
Microtubule (MT) nucleation and spindle bipolarisation were rescued by a point mutation in a γ-TuRC subunit or MT destabiliser. (A) (Left) Spindle bipolarisation after a prolonged monopolar state by a specific mutation in the *alp4^GCP2^* gene. Equatorial MTs during telophase were also recovered (arrow at 78 min). (Right) Failure in spindle bipolarisation. (B, C) Partial recovery of Alp6^GCP3^-GFP and MT intensities by a specific mutation in the *alp4^GCP2^* gene. The signal intensities from 12 to 15 min were compared between *plo1*Δ and *alp4-D440E plo1*Δ; Alp6^GCP3^-GFP intensity (**, p = 0.0049), and MT intensity (**, p = 0.0017). (D-G) Partial recovery of MT intensities by a mutation in the *asp1*^PPIP5K/Vip1^ gene. MT intensity from 12 to 15 min was compared between *plo1*Δ with *asp1-D507G plo1*Δ (****, p < 0.0001). In all the graphs, error bars indicate the standard error of the mean (SEM). Time 0 is set at the onset of spindle formation. Time 0 (min) is set at the onset of spindle formation.

Asp1^PPIP5K/Vip1^ is another MT-related factor and its mutation assisted in the growth of *plo1*Δ (Fig. 1B). Asp1^PPIP5K/Vip1^ is known to have a kinase domain at the N-terminus and a phosphatase domain at the C-terminus, and the latter is required for MT destabilisation (46). Interestingly, two mutations acquired during experimental evolution were located at the C-terminus (Fig. 1D). The mutation did not affect colony growth in the presence of Plo1 (Fig. 1C). However, time-lapse imaging showed that the *asp1-D507G plo1*Δ strain exhibited more spindle MT signals than *plo1*Δ, indicating that mutations in the C-terminal domain of Asp1 cause spindle MT stabilisation (Fig. 3D-G). These data suggested that bypass of Plo1 essentiality is achieved by increasing MT stability and/or generation.

### Glucose limitation bypasses Plo1 essentiality

The mechanism by which glucose limitation recovers the viability of *plo1*Δ is not readily explainable. Glucose reduction did not appear to change Plo1-GFP localisation. Both in high (3%) and low (0.08%) glucose media, Plo1-GFP was localised to SPBs from prophase to metaphase and delocalised at anaphase (Fig. S4A, B). To observe the process of mitosis, we followed Alp6^GCP3^-GFP and spindle MTs in double *ght5*Δ *plo1*Δ (Ght5 is a glucose transporter). Interestingly, Alp6^GCP3^-GFP accumulation at the SPB and spindle MT abundance were restored in the double mutant (Fig. 4A-D). MTs appeared to be more stable in *ght5*Δ, as incomplete disassembly of interphase MTs was often observed at the onset of mitosis, which reflected more total mCherry signals in the mutant than in the wild-type (arrows in Fig. 4B). Consistent with this observation, *ght5*Δ conferred resistance to TBZ (Fig. S3D).

**Figure 4.**
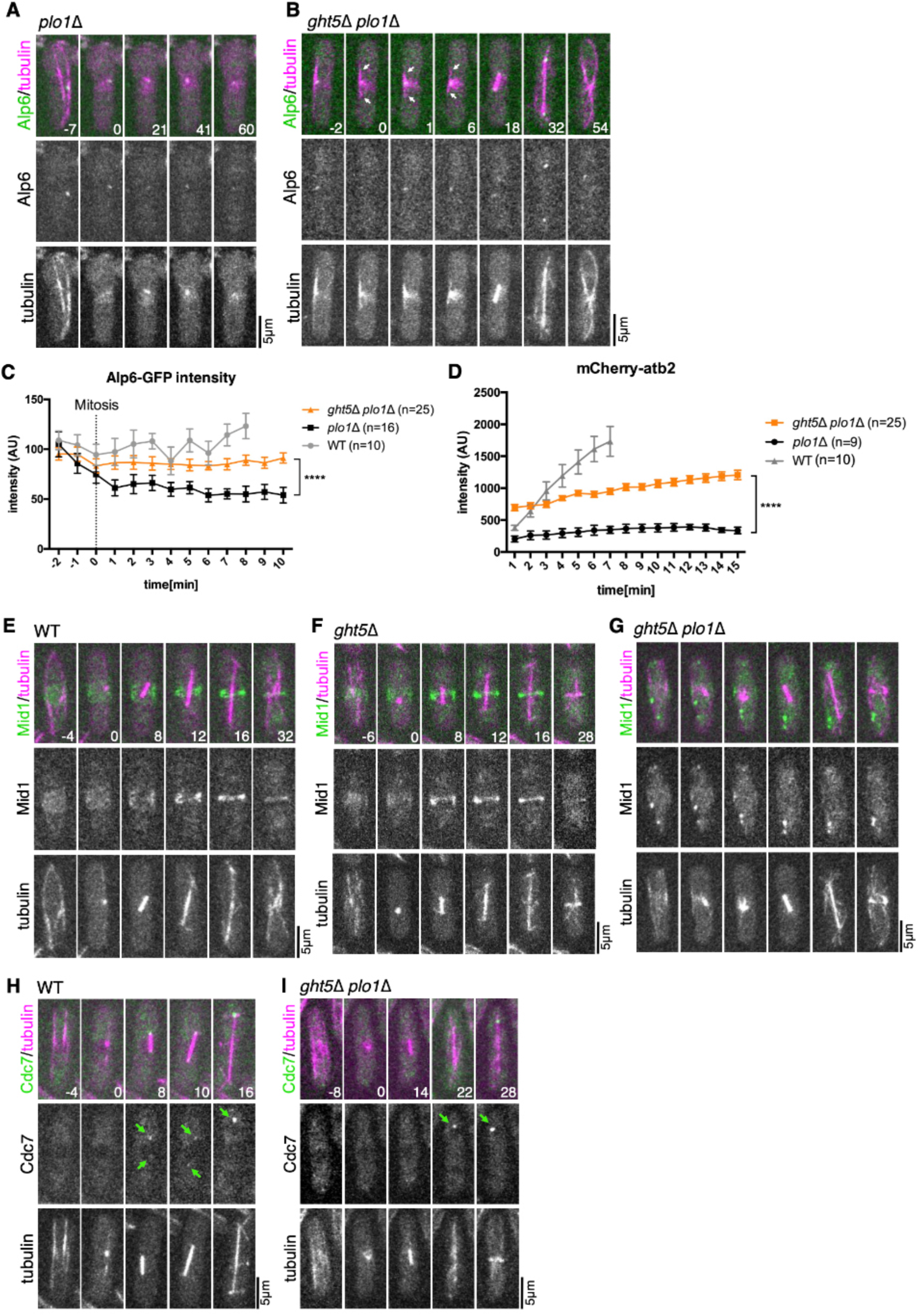
γ-TuRC localisation was restored by mutations in a glucose transporter in the absence of Plo1. (A) Live imaging of the *plo1*Δ strain expressing Alp6^GCP3^-GFP and mCherry-tubulin. The first mitotic phase after spore germination was imaged. (B) Live imaging of the *ght5*Δ *plo1*Δ strain expressing Alp6^GCP3^-GFP and mCherry-tubulin. Mitosis in the exponentially growing phase was imaged. Arrows indicate interphase microtubules (MTs) that remain during spindle assembly. (C, D) Quantification of Alp6^GCP3^-GFP and MT intensities during mitosis. The signal intensities were compared between *plo1*Δ with *ght5*Δ *plo1*Δ; Alp6^GCP3^-GFP intensity from 7 to 10 min (****, p < 0.0001) and MT intensity from 12 to 15 min (****, p < 0.0001). Error bars indicate the standard error of the mean (SEM). Time 0 is set at the onset of spindle formation. Control data is identical to that in Fig. 21 and J. The increase in MT intensity during the early mitotic stage in *ght5*Δ *plo1*Δ is due to the incomplete disassembly of interphase MTs (see arrows in B). (E-G) Equatorial accumulation of Midi ^annilin^-GFP is not restored in the viable *ght5*Δ *plo1*Δ strain. (H, I) Spindle pole body (SPB) localisation of Cdc7^Hippo^-GFP at metaphase is not restored in the viable *ght5*Δ *plo1*Δ strain. Arrows indicated GFP signals at SPBs. Time 0 (min) is set at the onset of spindle formation.

Next, we tested the localisation of Mid1^anillin^, which is recruited to the equatorial region during mitosis and defines the division site, depending on phosphorylation by Plo1 (47). We observed that Mid1 was not properly localised to the cortex in the viable *ght5*Δ *plo1*Δ strain, whereas the cortical localisation was normal in single *ght5*Δ (Fig. 4E-G). Consistent with this observation, the septum was mislocalised in *ght5*Δ *plo1*Δ, similar to *mid1*Δ (Fig. S4C-E). The results revealed that the septation error was not directly linked to the lethality of *plo1*Δ. Cdc7^Hippo^ is another downstream factor of Plo1; the SPB localisation of Cdc7^Hippo^ during metaphase, but not telophase, is impaired in the *plo1* mutant (48). In the viable *ght5*Δ *plo1*Δ strain, Cdc7^Hippo^-GFP localisation at metaphase SPB was not detectable (Fig. 4H, I). These results indicated that not all Plo1 downstream events, including the phosphorylation of the direct substrate, are restored by *ght5* mutations.

### Casein kinase 1 (CK1) constitutes a masked mechanism for spindle bipolarisation

Since proteins in the glucose-PKA pathway are not SPB-or spindle-associated, we hypothesised that other pathways are enhanced when glucose is limited, which promotes γ-TuRC localisation. To identify the effector proteins in such pathways, we performed a genetic screening, with the aim to acquire mutants that were synthetic lethal with double *ght5*Δ *plo1*Δ or *pka1*Δ *plo1*Δ. For this, we first transformed a plasmid containing the *plo1*^+^ gene in the double mutants, conducted mutagenesis, and selected the strains that could not lose the plasmid (Fig. 5A). A total of 13 mutants were identified that were synthetic lethal with either *ght5*Δ *plo1*Δ (seven strains) or *pka1*Δ *plo1*Δ (six strains). Possibly responsible genes were selected based on sequencing (e.g. dramatic amino acid changes, nonsense mutations, or identified in multiple strains). Synthetic lethality was confirmed for five genes (*bub1, hhp1, iml1, mak1*, and *wis1*) and one gene (*sin1*) by gene disruption and crossing with *ght5*Δ *plo1*Δ and *pka1*Δ *plo1*Δ, respectively. However, two mutants (*iml1* and *wis1*) and one mutant (*sin1*) resulted in poor growth when singly combined with *ght5*Δ and *pka1*Δ, respectively. These were excluded from further analysis because the major basis of synthetic lethality may not involve the lack of Plo1 kinase. *mak1* showed complex genetic interaction; while *mak1*Δ *ght5*Δ grew normally, synthetic lethality was revealed when mCherry-tubulin was introduced. In addition, the double *mak1*Δ *pka1*Δ grew poorly in the absence of mCherry-tubulin expression. Therefore, this gene was also excluded from further analyses. In contrast, triple disruption was not selected for two other genes, *bub1* and *hhp1* (Fig. S5A, B), whereas the double mutants with *ght5*Δ grew in a manner indistinguishable from the single *ght5*Δ even in the presence of mCherry-tubulin. We further confirmed the synthetic lethality of *hhp1*Δ with other PKA-pathway genes *git1*Δ *plo1*Δ and *pka1*Δ *plo1*Δ (Fig. S5C, D). Thus, *bub1* and *hhp1* were essential for *plo1*Δ viability.

**Figure 5.**
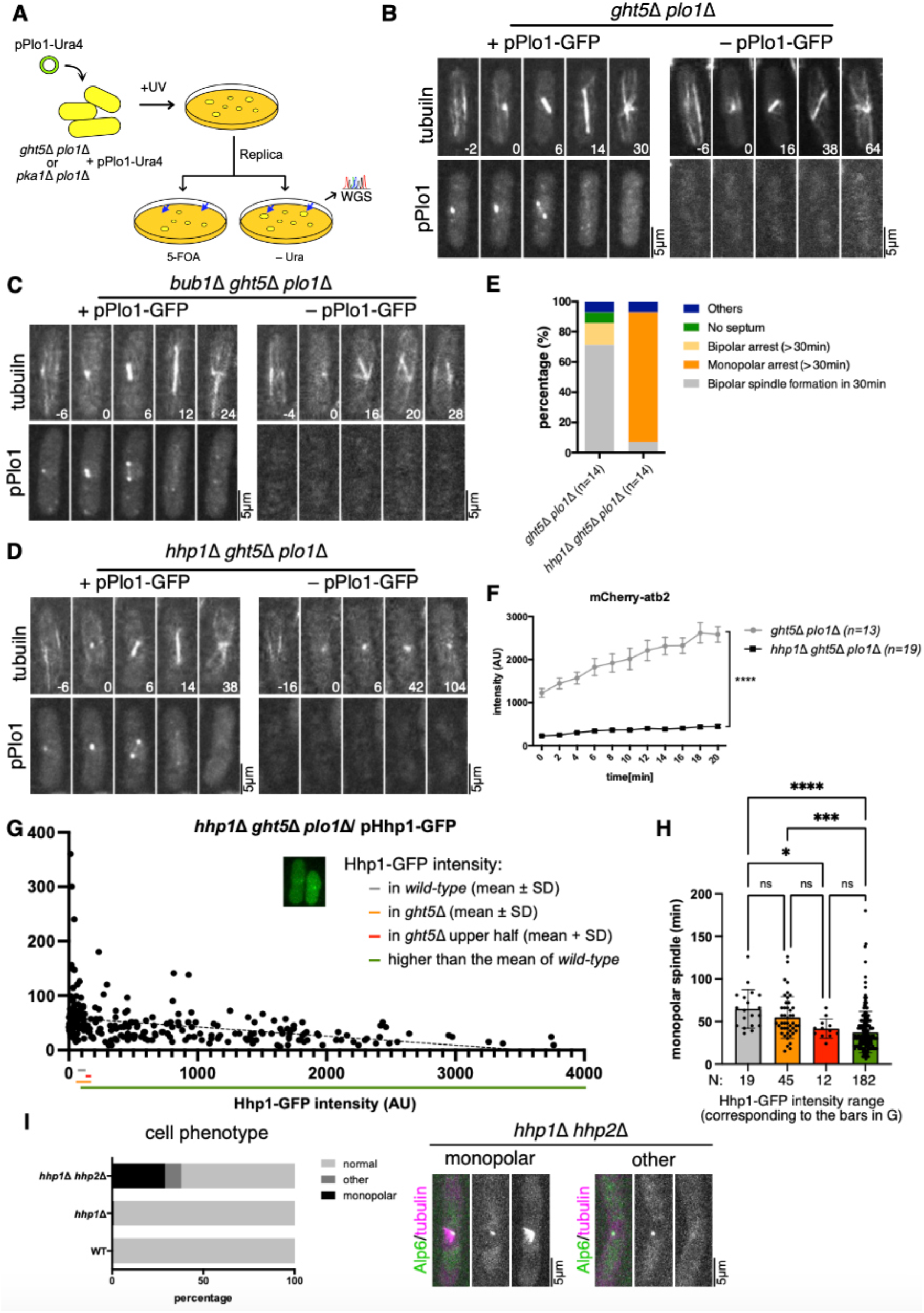
Hhp1^CK^) becomes essential for spindle formation in the absence of Plo1. (A) Schematic presentation of the synthetic lethal screening. The strain possessing Plo1 plasmid (*ura4+*) is sensitive to 5-FOA and therefore, docs not grow. The strain that cannot grow specifically on the 5-FOA plate should have a mutation that is synthetic lethal with *ght5*Δ *plo1*Δ or *pka1*Δ *plo1*Δ. The genome sequences of these strains were determined (‘WGS’). (B-D) Plasmid-loss experiment. The indicated double or triple disruptants transformed with Plo1-GFP plasmid were grown. Time-lapse imaging was performed and mitotic cells with or without Plo1-GFP signals were analysed. (E) Frequency of mitotic phenotypes (in the absence of Plo1-GFP). (F) MT intensity decreased in the absence of Hhp1^CKl^(****, p <0.0001). (G) Plasmid-loss experiment using a *plo1*Δ *ght5*Δ *hhp1*Δ triple disruptant and multicopy Hhp1-GFP plasmid (*lew*+). Time spent with monopolar spindles was plotted for each cell. GFP intensity (arbitrary unit) corresponds to the amount of Hhp1 in a cell. Mean intensities (± SD) of Hhp1-GFP signals in the wild-type background and *ght5*Δ background were indicated by grey and orange bars, respectively. In this analysis, the mean background intensity of the parental strain that had no GFP expression (80.3 AU, n = 31) was subtracted from the Hhp1-GFP intensity value. (H) Time required for monopolar-to-bipolar conversion. Three bars correspond to the samples with different GFP intensities described in (G). Error bars represent SD (*, p = 0.0458; ***, p =0.0001; ***, p <0.0001), and ns stands for “not significant.” (I) A total of 29% of the *hhp1*Δ *hhp2*Δ cells (n = 79) and 1% of the *hhp1*Δ cells (n = 272) assembled monopolar spindles, whereas this never occurred in control cells (n = 336). A lack of spindle MTs was also observed in the double disruptant (right). Time 0 (min) was set at the onset of spindle formation.

To identify the lethal event caused by these mutations, we observed live cells of the triple disruptants, *bub1*Δ *ght5*Δ *plo1*Δ and *hhp1*Δ *ght5*Δ *plo1*Δ. For this, we selected each triple disruptant that possessed the Plo1-GFP multicopy plasmid. Viable cells were cultured in non-selective medium, by which cells naturally lose the plasmid at a certain probability. Time-lapse images were then acquired. We analysed the cells that no longer had Plo1-GFP signals, as these cells represent triple gene disruptants. As a control, we prepared double *ght5*Δ *plo1*Δ possessing the Plo1-GFP plasmid and performed the identical ‘plasmid loss’ culture. In the control strain that had no GFP signals, monopolar spindles were converted into bipolar spindles within 30 min, followed by entry into anaphase, in >60% cells, as expected (Fig. 5B and E). In contrast, in triple *bub1*Δ *ght5*Δ *plo1*Δ, anaphase began even when spindles were still monopolar in 13 out of 43 cells (Fig. 5C). This phenotype explains the lethality of the strain and is consistent with the fact that Bub1 is an integral component of the spindle assembly checkpoint, which prevents premature anaphase entry (49). In contrast, in *hhp1*Δ *ght5*Δ *plo1*Δ, >80% of cells were arrested in monopolar states for >30 min, and spindle bipolarisation and anaphase entry were scarcely observed, similar to the *plo1*Δ strain in the normal medium (Fig. 5D and E). We concluded that the lethality of *hhp1*Δ *ght5*Δ *plo1*Δ comes from a defect in spindle bipolarisation, similar to *plo1*Δ in the normal medium.

*hhp1* encodes casein kinase I (CK1), which is distributed throughout the cell and is enriched at the SPB (50, 51). Hhp1^CK1^ is involved in a variety of cellular processes, such as DNA repair, ubiquitination-dependent regulation of septation initiation, DNA recombination and cohesin removal during meiosis (50, 52-54). However, to the best of our knowledge, Hhp1^CK1^ has not been directly linked to spindle function in fission yeast.

To investigate the basis of the unexpected genetic interaction, we first tested whether Hhp1 ^CK1^ expression/localisation was altered by *ght5* disruption. To this end, we tagged GFP to Hhp1 ^CK1^ in the wild-type and *ght5*Δ backgrounds. Time-lapse mitosis imaging and GFP intensity quantification indicated that Hhp1 ^CK1^ localisation was unchanged, but the overall abundance became more variable and on average slightly increased in the absence of *ght5* (96 ± 23, 120 ± 63 [AU, ± SD], n = 30 each). However, the level of *hhp1* mRNA was not elevated by a glucose reduction, suggesting that post-transcriptional regulation underlies the increased Hhp1 in the cell (Fig. S5E). Next, we tested whether the upregulation of Hhp1^CK1^ is necessary for the bypass of Plo1. We selected the *hhp1*Δ *ght5*Δ *plo1*Δ triple disruptunt that possesses the Hhp1^CK1^-GFP multicopy plasmid, and conducted a plasmid-loss experiment. In this experiment, GFP signal intensity served as an indicator of intracellular levels of the Hhp1^CK1^ protein. Time-lapse imaging and subsequent image analysis showed that the level of the Hhp1^CK1^ protein was overall correlated with the efficiency of spindle bipolarisation (Fig. 5G, 5H [grey vs. green]). However, the impact of the slight increase in Hhp1^CK1^ observed in *ght5*Δ was marginal; when we compared the time required for monopolar-to-bipolar conversion, we observed a slight and statistically nonsignificant decrease (Fig. 5H [grey vs. orange]). Thus, a moderate increase in Hhp1^CK1^ facilitates bipolar spindle formation in the absence of Plo1 and Ght5, although it may not be a prerequisite for BOE.

Next, we investigated whether ectopic expression of Hhp1 ^CK1^ was sufficient for the recovery of *plo1*Δ viability. We tested the expression of Hhp1^CK1^ by two different promoters on the multicopy plasmid, but we could not obtain data that reproducibly showed that Hhp1^CK1^ expression restored *plo1*Δ colonies (Fig. S5F). In addition, Hhp1^CK1^ expression from the plasmid did not enhance the growth of *alp6-V664F plo1*Δ or *alp4-D440E plo1*Δ, which was viable on its own but had slower growth than the wild-type (Fig. S5F). Thus, an increase in Hhp1^CK1^ levels alone did not increase the fitness of *plo1*Δ and was insufficient for the bypass of Plo1 essentiality.

Finally, we observed spindle dynamics in the *hhp1* single disruptant. Most of the cells (99%) assembled bipolar spindles, and mitosis proceeded comparably to the wild-type. However, among the 272 cells monitored, we found that 3 cells (1%) formed monopolar spindles; this was not observed in our imaging of control Hhp1^CK1^+ cells (N >336) (Fig. 5I). Furthermore, *hhp1*Δ *ght5*Δ was more sensitive to TBZ than *ght5*Δ (Fig. S3D). Thus, Hhp1^CK1^ has a very mild, almost negligible level of contribution to MT stability and bipolar spindle assembly in the presence of Plo1, but becomes essential in the absence of Plo1. In *S. pombe, hhp2*^+^ also encodes CK1 (52). Therefore, we selected the *hhp1*Δ *hhp2*Δ double disruptant expressing mCherry-tubulin and Alp6^GCP3^-GFP, and performed time-lapse microscopy. Interestingly, monopolar spindles appeared at a much higher frequency than single *hhp1*Δ (29%, n = 79) (Fig. 5I). Other phenotypes, such as undeveloped spindle MTs, were also observed in the double disruptant (Fig. 5I, right). We further determined if *hhp2*Δ would be synthetically lethal with three viable *plo1*Δ strains (*ght5*Δ *plo1*Δ, *git1*Δ *plo1*Δ, and *pka1*Δ *plo1*Δ). Unlike *hhp1*Δ, no strains showed synthetic lethality with *hhp2*Δ. Thus, Hhp1^CK1^ and Hhp2^CK1^ were not completely redundant for bypass-related functions, which corroborates the previous report that they are different in subcellular localisation and abundance (51).

### Masked contribution of CK1 to spindle formation in a human colon cancer cell line

Among the four Plks in mammals, Plk1 is required for centrosome maturation and bipolar spindle formation in many cell types and is thus most analogous to *S. pombe* Plo1. There are also several CK1 family members in mammals. As CK1δ is localised at the centrosome (55, 56), we tested whether CK1δ constitutes the masked mechanism behind Plk1 in human cells (Fig. 6). The treatment of a human colon cancer line (HCT116) with a low concentration (3 nM) of Plk1 inhibitor BI2536 resulted in a slightly higher frequency of monopolar spindle appearance in early prometaphase (Fig. 6A and B). PF670462, an inhibitor of CK1δ/ε (57, 58), did not increase the number of monopolar spindles. However, when both inhibitors were simultaneously treated, 36% of the cells first assembled monopolar spindles (Fig. 6A and B). The monopolar spindles were eventually converted to bipolar spindles; however, this process required >30 min in ~20% of the cells when two compounds were simultaneously added (Fig. 6C). These results highlight the importance of CK1, perhaps CK1δ, in spindle bipolarisation in human colon cancer cells, when Plk1 function is partially impaired.

**Figure 6.**
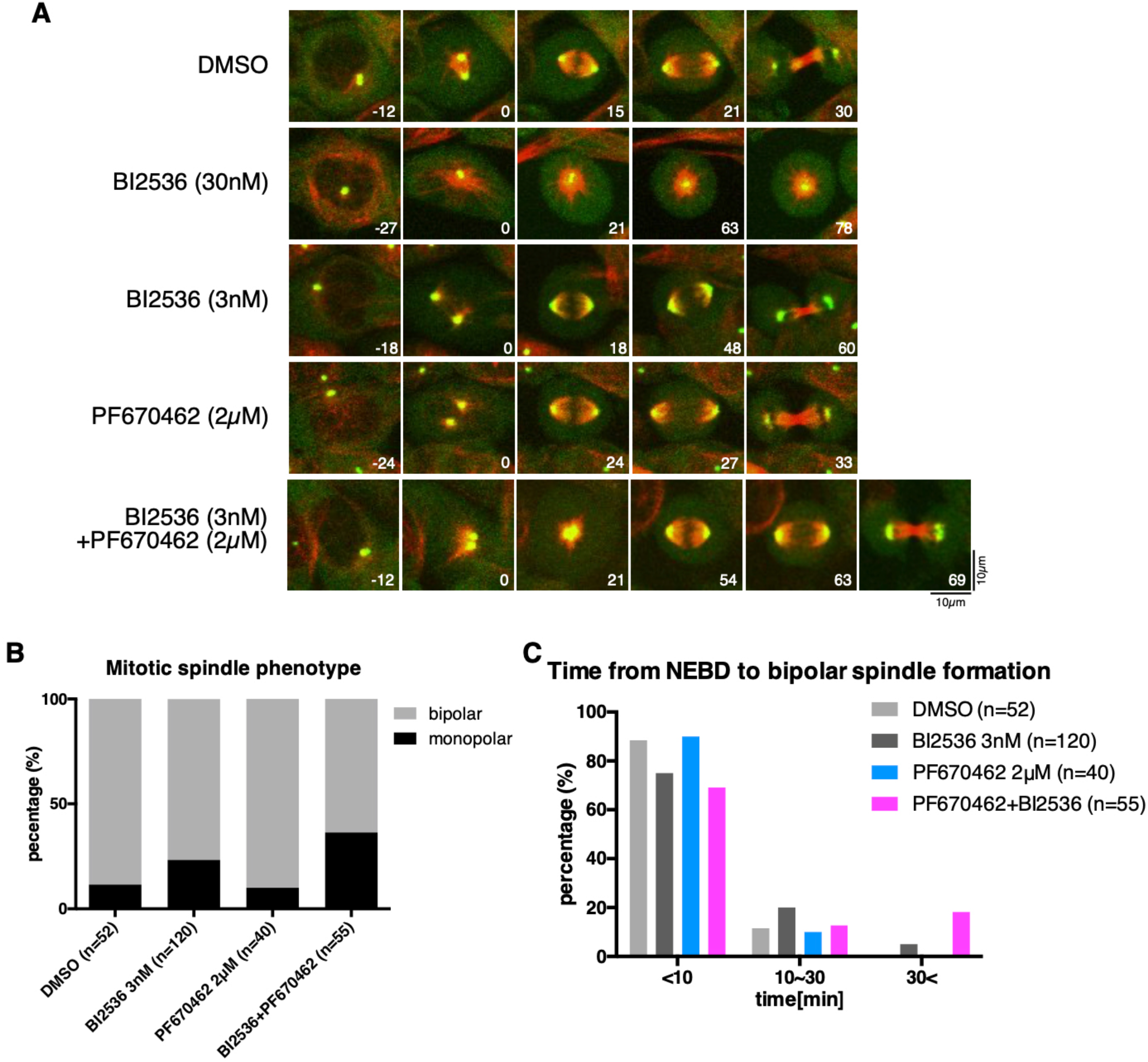
Synthetic monopolar spindle phenotype by partial inhibition of Polo-like kinase (Plk)1 and casein kinase I (CKl)δ in human colon cancer cells. (A) Mitosis of the HCT116 cell line in the presence of Plk1 and/or CK1 inhibitors (BĪ2536 for Plk1, PF670462 for CK1). Green, γ-tubulin-mClover (endogenously tagged); Red, Sir-tubulin. (B) Frequency of monopolar spindles (monopolar state for ≥10 min). (C) Duration of nuclear envelope breakdown (NEBD)-to-bipolar spindle formation. Time 0 (min) is set at the onset of spindle formation.

## Discussion

This study represents a rare example of the experimental BOE of genes required for mitosis. The BOE occurrence in Plo1 was unexpected, as it has been recognised as a versatile, essential kinase in mitosis not only in animal cells but also in fission yeast. However, there is evolutionary evidence supporting that this gene can be deletable; for example, plants have lost Plks, whereas the ancestral algae possess Plks (59). In our initial BOE screening using *plo1*Δ spores, only one viable strain was recovered, in which the gene encoding the glucose transporter Ght5 was lost through a deletion event. Subsequent evolutionary repair (experimental evolution) experiments led to the identification of more mutations, many of which restored viability of *plo1*Δ without the *ght5* mutation. Thus, the initial mutagenesis-based screen was not sensitive enough to capture all the possible BOE. More BOE may be uncovered in the yeast system, including BOE-M, by applying more sensitive methods, or simply by increasing the screen scale.

Plo1 loss can be rendered non-lethal by mutations in several genes, some of which were unrelated to each other and not associated with spindle functions at first glance. However, this is in accordance with many previous examples of BOE or evolutionary repair in the laboratory, where compensatory mutations are often found in genes outside of the perturbed functional module (3). Subsequent analysis suggested that bypass mutations converge into a common outcome: the increase in spindle MTs. This was achieved by multiple direct and indirect mechanisms, such as mutations in an MT destabiliser and MT nucleator, or through glucose starvation. In contrast, the septum phenotype was not rescued in a viable strain. Thus, although multiple defects have been identified in the *plo1* mutants, MT formation is directly linked to viability. In a broader sense, BOE analysis could be used to distinguish between essential and non-essential functions of an essential gene.

The bypass of Plo1 essentiality by glucose reduction in the medium is intriguing from multiple perspectives. First, it illustrates the non-absolute nature of gene essentiality (1). If the low-glucose medium was used as the standard yeast culture medium, then Plo1 would have been assigned as a non-essential gene in *S. pombe*. Second, a change in available nutrients occurs, perhaps frequently, in the natural yeast habitat. The decrease in available glucose allows the yeast to lose a critical mitotic kinase and develop an alternative mechanism. The data support the theory that environmental change combined with gene mutations drives molecular diversity, namely, variation in genes required for an essential process (3). Third, the change in fundamental metabolism alters the expression of many genes (60), offering a unique ‘genetic background’ that is not achieved by mutations in a few mitotic genes. In the case of *plo1*Δ, a critical factor for survival was Hhp1^CK1^. Because Hhp1^CK1^ is SPB-associated, it is possible that critical Plo1 substrates (such as γ-TuRC or its associated factor) are phosphorylated by Hhp1^CK1^. However, CK1 is unlikely the sole element of BOE based on glucose repression and other factors should be also involved, as Hhp1 ^CK1^ overexpression alone was not sufficient to restore the viability of *plo1*Δ (Fig. 7). Interestingly, *S. cerevisiae* Hrr25^CK1^ can phosphorylate and activate the γ-tubulin complex *in vitro* and this phosphorylation is required for *in vivo* γ-tubulin functions (61). Whether Hrr25^CK1^ constitutes a masked mechanism of Cdc5^Plk1^ in *S. cerevisiae* is an intriguing question for future investigation.

**Figure 7:**
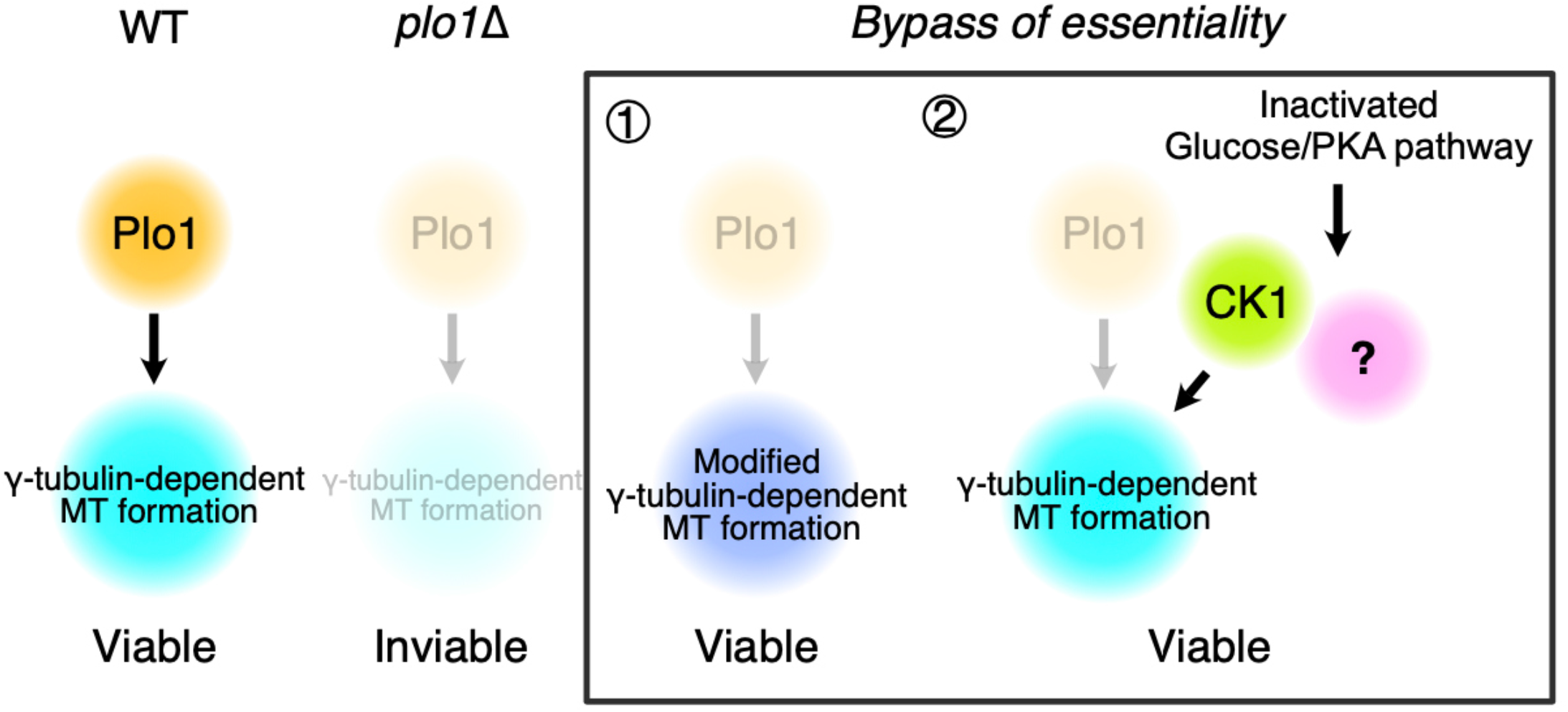
How Plo1 essentiality is bypassed. An increase in spindle microtubules (MTs) is the key to bypassing Plo1. This can be achieved by mutations in MT-associated proteins or nucleators (1) or global change in glucose metabolism (2), which involves casein kinase I and other unknown factors.

BOE, or synthetic viability, is a critical challenge in cancer chemotherapy because of the emergence of resistance (62). Plk inhibitors have been recognised as promising antitumor drugs (21, 23). However, our study suggests that there may be resistant cells involving CK1 and that double inhibition of Plk1 and CK1δ may be more suitable for mitotic cell perturbation.

## Materials and methods

### Yeast strains and media

We followed Takeda et al. (2019) for the yeast culture and gene disruption. Complete medium YE5S contained 1% yeast extract and 3% (w/v) glucose, supplemented with adenine, leucine, histidine, uracil, and lysine, whereas glucose was reduced to 0.02% in the low-glucose medium. Synthetic PMG and EMM media were used when selection was based on adenine, leucine or uracil. Cells were cultured at 32°C (plate) or 30°C (liquid), unless other temperatures are indicated. Sporulation was induced in an SPA plate. Gene disruption, site-directed mutagenesis, and GFP/mCherry tagging were performed using the standard one-step replacement method (i.e. homologous recombination). Transformation was performed using the conventional LiAc/PEG method (63), and targeted integration was confirmed by PCR. Selection of the strain after random spore or transformation was based on leucine, uracil, G418 (100 μg/mL), hygromycin (50 μg/mL), clonNAT (100 μg/mL), or blasticidin S deaminase (37.5 μg/mL). The strains, plasmids, and primers used in this study are listed in Tables S1, S2, and S3, respectively.

### Human cell culture

The human colon cancer-derived HCT116 line, in which the endogenous TubG1 gene (γ-tubulin) was tagged with mClover, was cultured in McCoy’s 5A medium (Gibco) supplemented with 10% serum and 1% antibiotics (64). Plk1 inhibitor BI2536 (3 nM) and CK1 inhibitor PF670462 (2 μM) were treated for 24-30 h prior to imaging, whereas 30 nM BI2536 was added at the beginning of imaging. MTs were stained with 50 nM SiR-tubulin for 2 h prior to imaging; this concentration of SiR-tubulin did not significantly affect MT growth and nucleation in this cell line (64).

### BOE screening and confirmation

We followed the method described by Takeda et al. (2019). The initial screening encompassing eight mitotic gene disruptants was performed for 1 × 10^7^ spores, and *plo1* screening was repeated with a larger scale (8 × 10^7^ spores). One or 11 viable colonies were obtained for *plo1* in the respective screening, for which PCR confirmed that the strain was indeed deleted from the entire *plo1* open reading frame (ORF). One of the viable *plo1*Δ strains was crossed with the wild-type (975 or 972), followed by sporulation on an SPA plate. The spores were incubated in YE5S with G418 (100 μg/mL) and cycloheximide (100 μg/mL) plates. The viable colonies were subjected to WGS (i.e. bulk segregant analysis). Analysis of the genome sequence of the strain identified a unique ~7 kb deletion on chromosome III, in which three genes, *SPCC1235.17, SPCC1235.18*, and *ght5*, were included. By crossing other lines with the Ght5-GFP integrant strain (24), we observed a strong genetic link for six of the strains (one strain was sterile and was not analysed further). Sequencing of the three strains verified that all had mutations or deletions in the *ght5* locus; one showed a mutation from TGG to TAG that introduced a premature stop codon in *ght5* and another showed Gln^152^ to Pro substitution of *ght5*. To confirm this suppression, the *ght5* gene was independently deleted by homologous recombination, and the growth of *ght5*Δ *plo1*Δ cells was investigated using a spot test. WGS of the other three *plo1*Δ revertants identified other suppressor candidates (the other line was sterile and was not further analysed). One strain had TT instead of AC in the *ngg1* coding region, which introduces a premature stop codon. Ngg1 (also called Ada3) is a component of the SAGA complex (65). To investigate whether the mutations in *ngg1* rescue *plo1* lethality, a genetic linkage test was conducted with the *rec6* gene, which is located close to *ngg1*. The other two revertants had mutations in the *cyr1* gene (encoding adenylate cyclase, a component of the PKA pathway). Nucleotide insertion in *cyr1* of one strain caused a frameshift that induced a premature stop codon, whereas the other strain had an amino acid change from Asn to Lys. Therefore, the *plo1*Δ rescue ability by *pka1* or *cyr1* deletion was tested. The strains lacking both *pka1* and *plo1* or *cyr1* and *plo1* grew on 3% glucose medium, albeit slower than the wild-type.

### Whole genome sequencing

Approximately 2 × 10^8^ cells were harvested, and their genomic DNA was extracted using Dr. GenTLE (Takara). The genomic DNA was sequenced by BGI, Novogene, or the gene sequencing facility at Nagoya University, where the read length of the sequence was 150 bp or 81 bp. The FASTQ file was modified and mapped using CLC genomic workbench software. The ends (5 bp from the 5’ end and 1 bp from the 3’ end) of each fragment were trimmed, and these fragments were then filtered for a minimum sequencing quality of 30 datasets to reduce the error ratio. The fragments were aligned to the *S. pombe* genome reference and formatted as BAM files. IGV was used to visualise the BAM file and check the mutation site. Insertion, deletion, single nucleotide variation (SNV), and large indels were investigated. All the WGS data are available at Sequence Read Archive (SRA) at NCBI (accession number: PRJNA768628).

### Experimental evolution

Experimental evolution was conducted with serial dilution and saturation (66). Saturated culture after passage (10 mL) was 1,000-fold diluted for the next passage. After culturing for 150-200 generations, cell cultures were spread onto agar plates, and the fastest growing colonies were selected for each culture. In the first round, six *plo1*Δ colonies were inoculated into 0.08% glucose liquid medium with G418 (10 μg/mL; in order to avoid microbial contamination) and cultured at 30°C. Eventually, six independent colonies were picked up (hereafter called ‘1 ^st^ EVO’ strains). They showed faster colony growth on 0.08% glucose agar plate than the original *plo1*Δ strain. Three of them also formed small colonies on the normal 3% glucose medium. These strains were further subjected to experimental evolution using 3% glucose medium with G418 (10 μg/mL) at 30°C (two or three independent cultures). In the end, eight ‘evolved’ strains (i.e. the fastest growing strains) were obtained as *plo1*Δ second-round evolution strains (hereafter called 2^nd^ EVO). Another round of experimental evolution (3^rd^ EVO) was carried out for *gitl*Δ *plo1*Δ and *pka1*Δ *plo1*Δ strains at 36°C (two and six independent cultures, respectively), as these formed tiny colonies at 36°C, whereas the *plo1*Δ strain did not grow even in the low-glucose medium at this high temperature. We determined the genome sequences and identified the specific mutations in five out of six strains from 1 ^st^ EVO, including three strains for which 2^nd^ EVO selection was carried out, and six out of eight strains from 2^nd^ EVO. On average, we found 2.2 mutations introduced in the open reading frame. Three out of the five 1 ^st^ EVO strains had a point mutation in SAGA complex genes (*spt20, sgf73*, and *spt7*) Mutations in glucose transporters or PKA-pathway genes (*ght5, git5, git1*, and *ght8*) were found in four of the six 2^nd^ EVO strains. From the 3^rd^ EVO, we sequenced the three fastest growing clones and found *asp1* mutations in two strains and *tra1* (SAGA) in the third strain.

### Synthetic lethality screening

Double disruptants *ght5*Δ *plo1*Δ and *pka1*Δ *plo1*Δ were transformed with the Plo1 plasmid (*ura4*+) Cells (5 × 10^4^ - 1 × 10^6^ cells) were mutagenised with UV (150 or 200 × 100 μJ/CM^2^; UVP Crosslinker CL-1000) and plated onto a PMG (*ura4*-) plate. After 6 d, the colonies were replica plated onto YE5S plates. After 2 d, the colonies were replica plated onto 5-FOA and PMG (*ura4*-) plates. Cells that could not survive in the absence of the plasmid were isolated (inviable colonies after replica plating onto 2 mg/mL 5-FOA-containing plates). For one of the obtained strains, the genomic DNA library (pTN-L1, NBRP) was transformed into the strain, and the colonies found on 5-FOA plates were isolated. The plasmid extracted from the colonies was sequenced and found to encode *hhp1*^+^. However, this method did not work well for other strains. Therefore, for the other 12 strains, whole genome sequences were determined and candidate mutations (nonsynonymous and frameshift) were inspected by independent deletion and crossing.

### Microscopy

Exponentially growing yeast cells or spores were attached to a 1 μg/mL lectin-coated glass plate for >10 min at 32°C (67). Live imaging was performed with a spinning-disc confocal microscope (Nikon Ti; 100× 1.45 NA) with a perfect focus system. The cells were maintained at 32°C in a stage-top heater. Images were taken using an ImagEM CCD camera (Hamamatsu) every 1 −3 min for exponentially growing cells or every 1 or 2 min for germinating spores (z-stacks: 1 μm × 5 sections). Spores were pre-incubated for 10–12 h in 3% YE5S medium before imaging. For imaging that involved plasmid loss, cells were pre-incubated overnight in PMG without uracil or leucine and exponentially growing cells were transferred to 3% YE5S medium for ~24 h prior to microscopy. Human cell live imaging was performed with a spinning-disc confocal microscope (Nikon Ti; 40× 1.45 NA) with a perfect focus system. Images were captured using an ImagEM CCD camera (Hamamatsu). The cells were maintained at 37°C in a stage-top heater, where CO_2_ was supplied. Z-stack images (3 μm × 5 sections) were taken every 3 min. Images were analysed using FIJI, and the data were plotted using GraphPad Prism software. The signal intensity of mCherry-tubulin on the spindle and Hhp1-GFP in late G2 was measured after maximum projection, whereas a single focal plane was selected for Alp6-GFP and Cut7-GFP measurements. Two Alp6-GFP signals on the bipolar spindles were summed to obtain the total intensity. The maximum projection images are presented in the figures.

### Real time PCR

Exponentially growing yeast cells were lysed with zymolyase (50 U per 1 × 10^7^ cells), and total RNA samples were prepared using the RNeasy Plant Mini kit (Qiagen), using the yeast protocol described in the manufacturer’s instructions of RNeasy Mini Kit (Qiagen). To eliminate genomic DNA contamination, an additional DNase treatment was performed using an RNase-free DNase Set (Qiagen). PrimeScript II (Takara) was used for the reverse transcription. Real-time PCR was performed and analysed using Step One Plus (Applied Biosystems) with SYBR Green Master Mix (Applied Biosystems). The primer set designed at the C-terminus of *act1* was used as the control (24). The copy number of *hhp1* mRNA relative to *act1* was calculated from a standard curve drawn using serial dilutions of cDNA as the templates.

### Statics

All statistical analyses were performed using the GraphPad Prism software. Two-way ANOVAs were applied between the two groups in Fig. 3B, 3C, 3G, 4C, 4D, and 5F, and a two-way ANOVA with multiple comparisons using Tukey’s test to compare the four groups in Fig. 2H and a one-way ANOVA with multiple comparisons using Tukey’s test to compare the four groups (Fig. 5H). The data distribution was assumed to be normal, but this was not formally tested. Established P values were denoted as follows: P > 0.05 (ns), P < 0.05 (*), P < 0.01 (**), P < 0.001 (***), and P < 0.0001 (****). Error bars in the graph represent the SEM or SD of each group. Experiments were performed twice or more, and the data from one experimental set were presented after quantitative analysis, except for Fig. 4C, 4D, 5G, 5H, and 6C, where the data from multiple experiments were combined because of insufficient sample numbers in a single experiment.

## Acknowledgements

We are grateful to Aoi Takeda for help with the experimental evolution culture; Shigeaki Saitoh (Kurume University), Kojiro Takeda (Konan University), Masamitsu Sato (Waseda University), Iain Hagan (University of Manchester), Ye Dee Tay (University of Edinburgh), and National Bio-Resource Project (NBRP) of the MEXT, Japan, for yeast strains and plasmids; Kazuma Uesaka for help with sequence analysis; Ken Sawin, Hiro Ohkura, and Ye Dee Tay (University of Edinburgh) for valuable discussions and protocols; Rie Inaba, Kyoko Zenbutsu, and Miki Ueda for media preparation; Shigeaki Saitoh and Moé Yamada for comments on the manuscript. This work was supported by JSPS KAKENHI (19K22383) to G.G. and JST SPRING to J.K. The authors declare no conflicts of interest.

---

**Figure S1.**
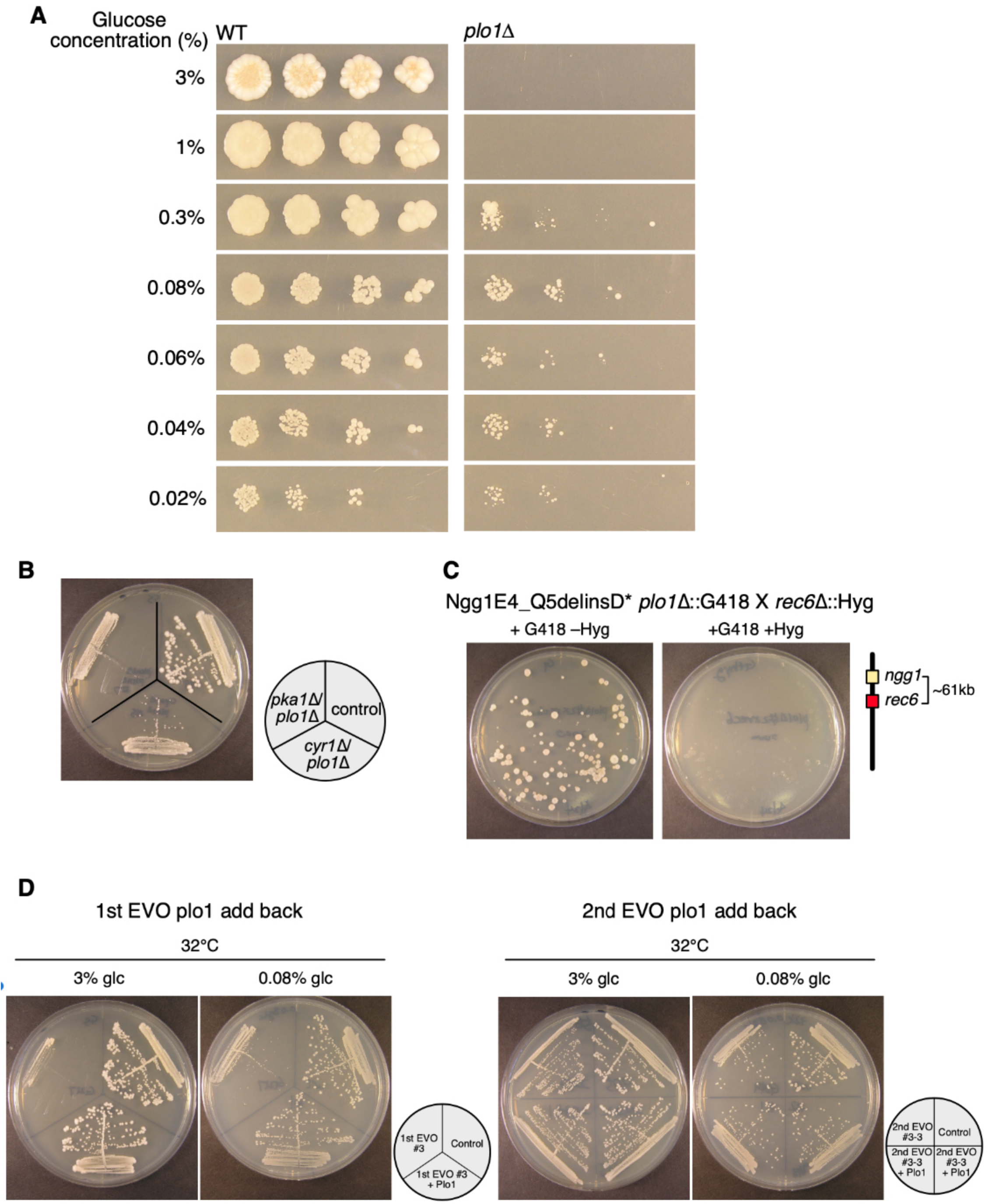
The bypass of essentiality (BOE) of Plo1. (A) Colony spotting on YE5S (+ G418, CHX) plates with various glucose concentrations. Spores (7,500, 1,500, 300, or 60) were spotted and incubated for 5 d at 32°C (expected drug-resistant cell numbers: 1,875, 375, 75, and 15, respectively). (B) Disruptant of PKA (*pka1*) or adenylate cyclase (*cyr1*) bypassed Plo1 essentiality. Colonics were grown on YE5S plates for 4 d at 32°C. (C) An example of suppressor mutation confirmation based on linkage test, *ngg1* and *rec6* genes are closely located on chromosome II. Double *plo1*Δ *rec6*Δ was never obtained after crossing *rec6*Δ (marked with hygromycin resistance) with a surviving *plo1*Δ strain (G418 resistant) in which a mutation was identified in *ngg1*. (D) Experimental evolution (EVO) of *plo1*Δ strain was performed sequentially, as described in Fig. 1 A. After each round, we isolated a few of the fastest growing colonies. We converted *plo1*Δ to *plo1*+ and tested if this enhanced the colony growth. Return of Plo1 helped colony growth of a strain obtained after the 1^st^ round of evolution (left, strain #3), whereas a further evolved strain was insensitive to Plo1 (right, strain #3-3). Two glucose concentrations were tested (3% and 0.08%). Wild-type was used as the control.

**Figure S2.**
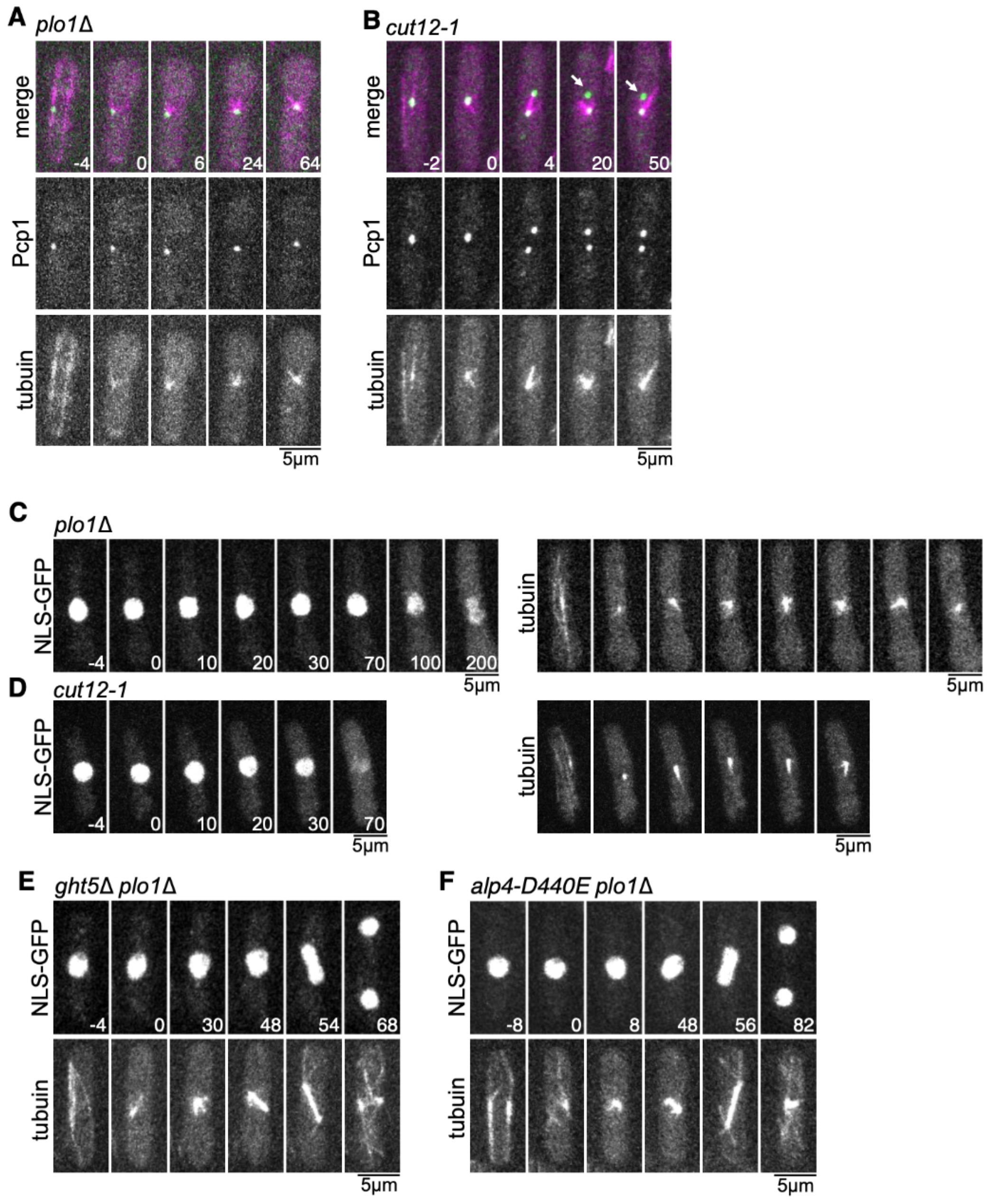
Nuclear envelope appears to be intact in *plo1*Δ. (A, B) Live imaging of *plo1*Δ and *cut12-1* strains expressing Pcp1^pericentrin^-GFP and mCherry-tubulin. *cut12-1* cells were incubated at 36°C for 3 h before imaging. (C-F) Live imaging of *plo1*Δ, *cutl2-1, ght5*Δ *plo1*Δ, and *alp4-D440E plo1*Δ strains expressing NLS-GFP-β-Gal and mCherry-tubulin. *cut12-1* cells were incubated at 36°C for 3 h before imaging. Time 0 (min) was set at the onset of spindle formation.

**Figure S3.**
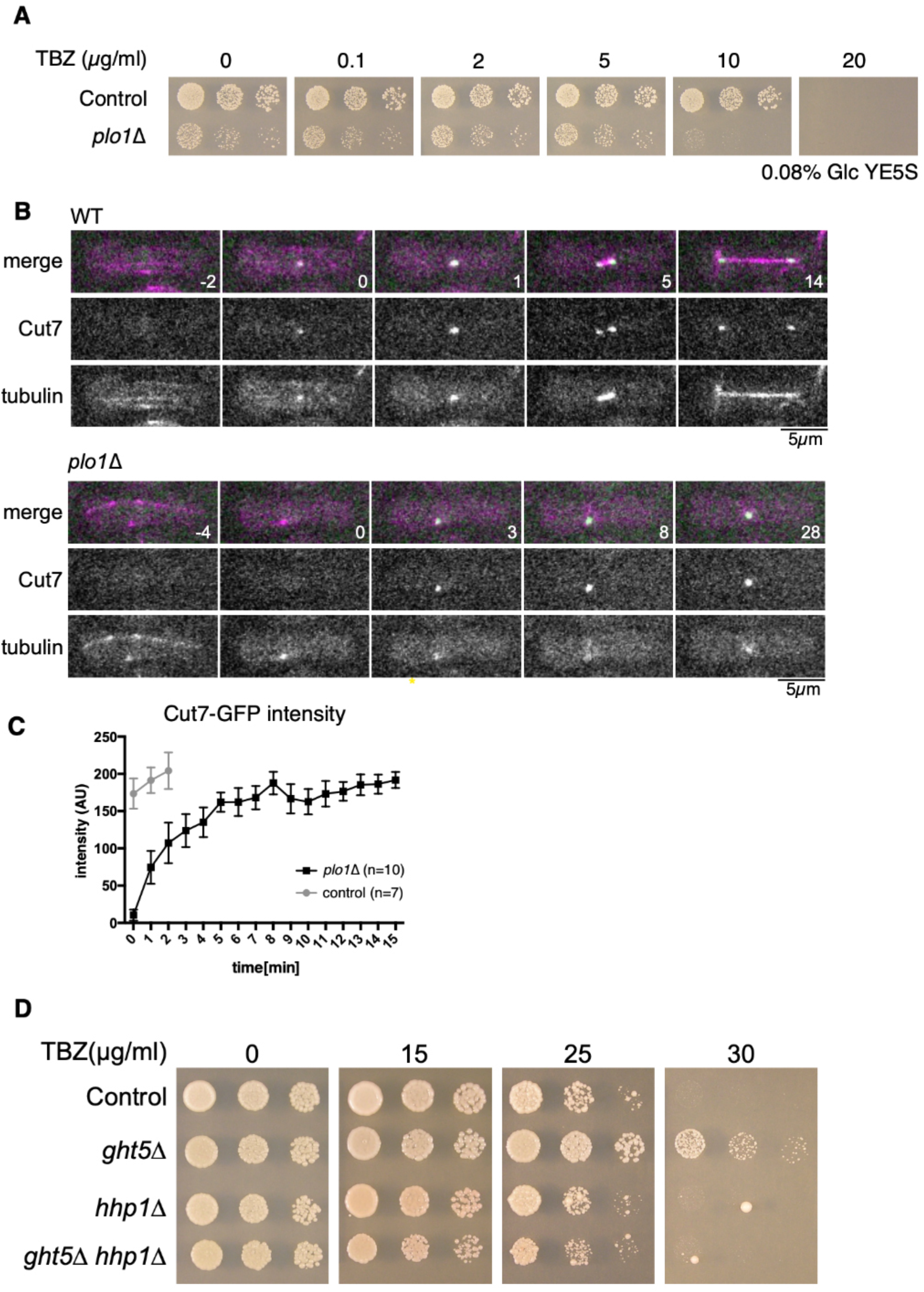
*plo1* deletion is sensitive to thiabendazole (TBZ) but does not critically affect Cut7^kinesin-5^ localisation. (A) In the presence of 10 μg/mL TBZ, *plo1*Δ failed to form colonies even in low glucose medium. Spores (10,000, 2,000, and 400) were spotted and colonies were formed on 0.08% glucose, G418-containing plates for 5 d at 32°C (expected G418-resistant cell numbers: 5,000, 1,000, and 200 cells, respectively). (B, C) Failure in spindle bipolarisation even when Cut7^kinesin-5^-GFP accumulated on the SPB at a normal level after a delay in *plo1*Δ. (D) *ght5*Δ confers resistance to TBZ, whereas *hhp1*Δ suppresses the effect. Cells (5,000, 1,000, and 200 cells) spotted and cultured in YE5S medium for 3 d (0 and 15 μg/mL TBZ plates) or 6 d (25 and 30 μg/mL TBZ plates) at 32°C. Time 0 (min) is set at the onset of spindle formation.

**Figure S4.**
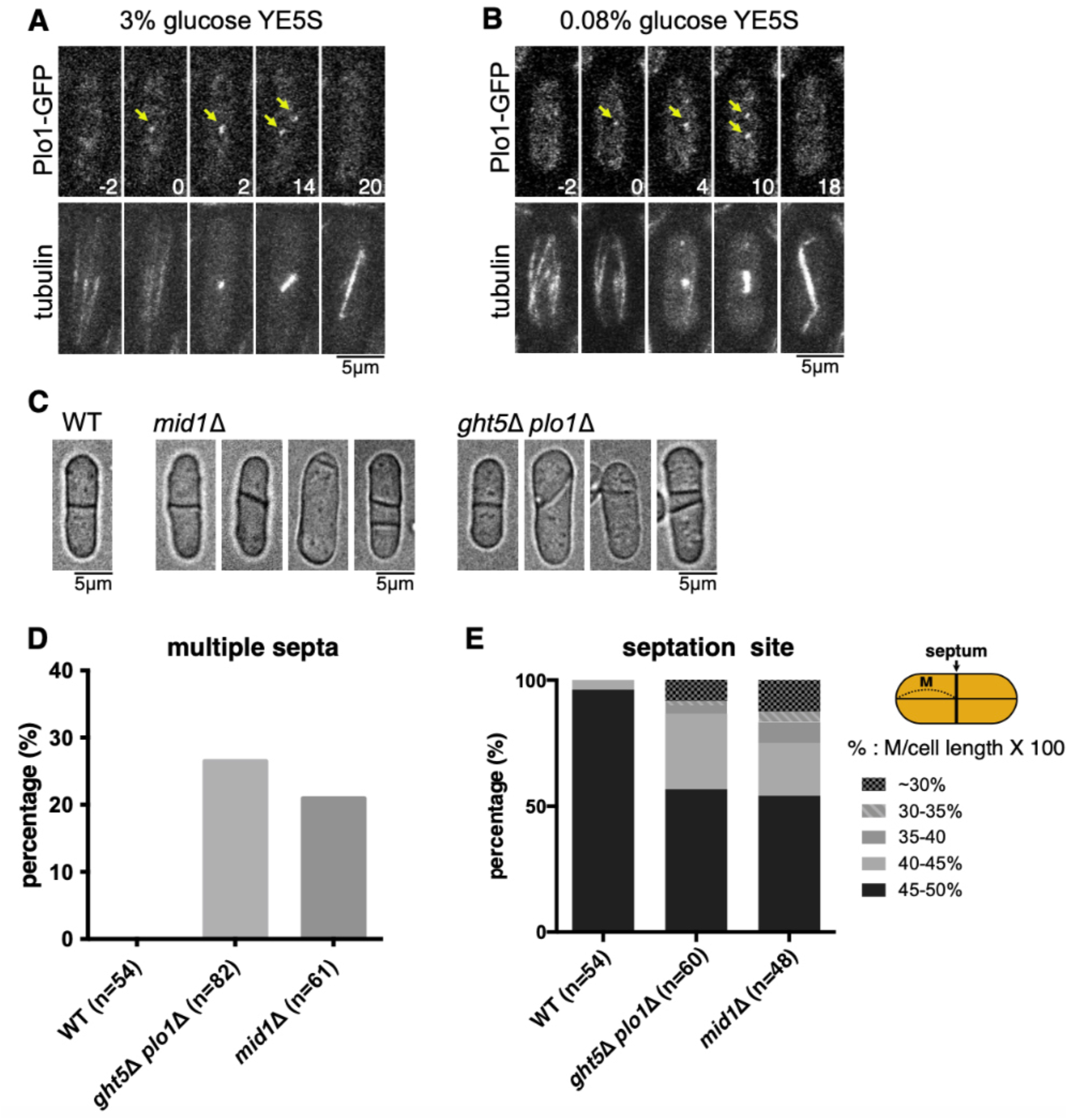
Septation defect of *plo1*Δ is not rescued by *ght5* deletion. (A, B) Live imaging of *plo1*Δ strains expressing Plo1-GFP and mCherry-tubulin. Cells were incubated in the YE5S medium containing 3% or 0.08% glucose. Arrows indicate Plo1-GFP signals at SPBs. Time 0 (min) was set at the onset of spindle formation. (C) Bright field images of wild-type (WT), *mid1*Δ, and *ght5*Δ *plo1*Δ. (D) Quantification of multiseptated cells. A total of 21% of *mid1*Δ cells and 26% of *ght5*Δ *plo1*Δ cells had multiple septations, whereas this was never observed in control cells. (E) Quantification of the septation site. Relative position of the septum along the long axis of the cell was determined.

**Figure S5.**
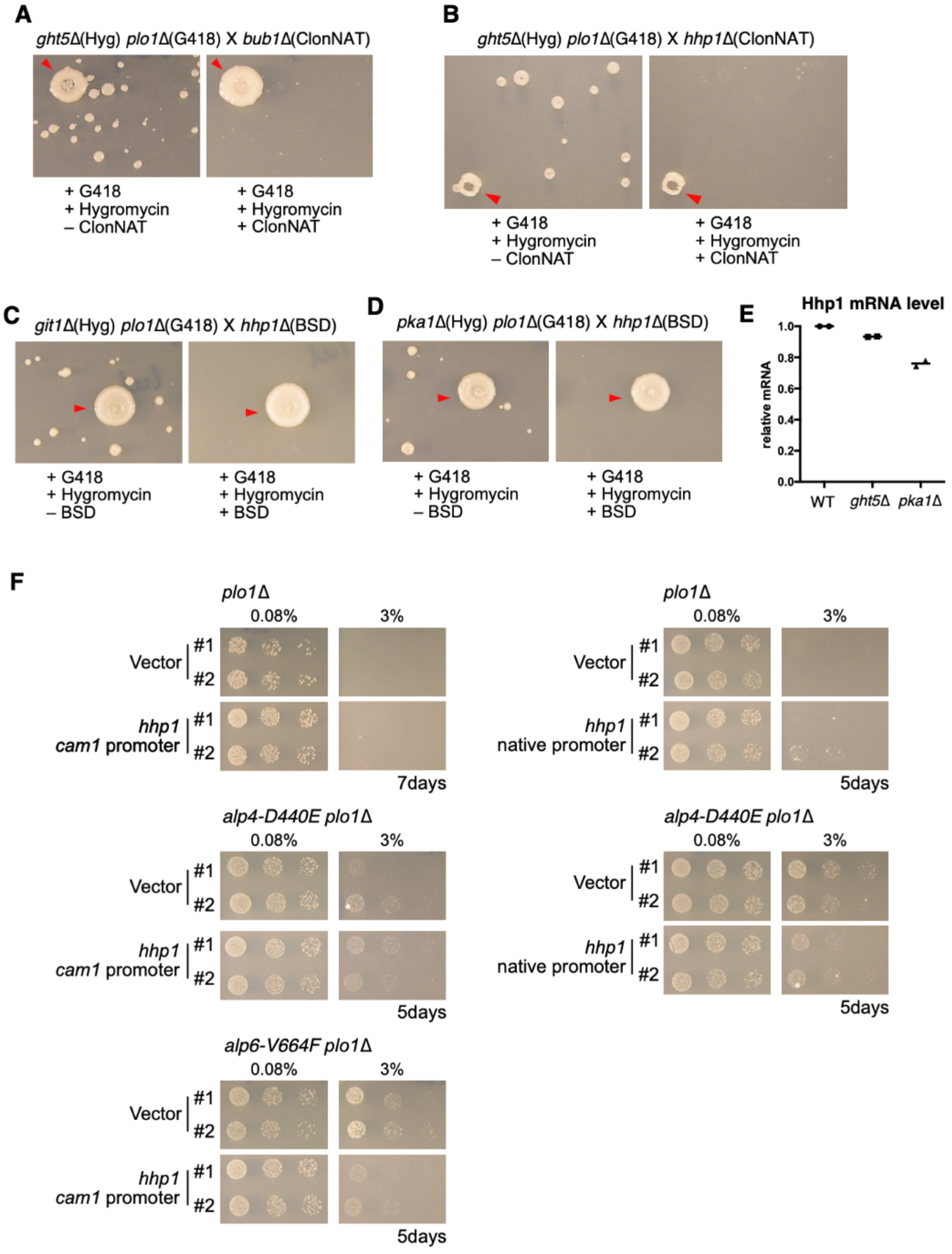
*ght5*Δ *plo1*Δ is synthetic lethal with *hhp1*Δ, but Hhp1^CK1^ overexpression alone does not increase the colony growth of *plo1*Δ. (A-D) Confirmation of synthetic lethality between the indicated strains by random spore spreading followed by replica-plating onto drugcontaining plates. The *hub1* and *hhp1* ORFs were replaced with ClonNAT-resistant or BSD-resistant cassettes. The *plo1* ORF was replaced with a G418-resistant cassette, *ght5, git1*, and *pka1* were replaced with a hygromycin-resistant cassette. Colony formation of heterozygous diploids (red arrowheads) indicates the functionality of the plate. (E) Relative *hhp1* mRNA levels in wild-type (WT), *ght5*Δ, and *pka1*Δ. Data from two independent experiments were plotted. (F) Investigation of the rescue ability by Hhp1 overexpression in the indicated strains. The calmodulin promoter (*caml-pr*) (67) and the native promoter of *hhp1*^+^ gene were used. Hhp1 was expressed in the *plo1*Δ, *alp4-D440E plo1*Δ, and *alp6-V664F plo1*Δ strains. Cells (10,000, 5,000, and 1,000) were spotted and cultured in YE5S with G418 for 5 d at 32°C.

